# Rhythms of the Body, Rhythms of the Brain: Respiration, Neural Oscillations, and Embodied Cognition

**DOI:** 10.1101/174276

**Authors:** Somogy Varga, Detlef H. Heck

**Affiliations:** Dept. of Philosophy, University of Memphis, Memphis, TN 38152; Dept. of Anatomy and Neurobiology, University of Tennessee Health Science Center, Memphis, TN 38163

**Keywords:** Neural oscillation, coherence, synchrony, embodied cognition, sensory afference, respiration, gesture

## Abstract

In spite of its importance as a life-defining rhythmic movement and its constant rhythmic contraction and relaxation of the body, respiration has not received attention in Embodied Cognition (EC) literature. Our paper aims to show that (1) respiration exerts significant and unexpected bottom-up influence on cognitive processes, and (2) it does so by modulating neural synchronization that underlies specific cognitive processes. Then, (3) we suggest that the particular example of respiration may function as a model for a general mechanism through which the body influences cognitive functioning. Finally, (4) we work out the implications for embodied cognition, draw a parallel to the role of gesture, and argue that respiration sometimes plays a double, pragmatic and epistemic, role, which reduces the cognitive load. In such cases, consistent with EC, the overall cognitive activity includes a loop-like interaction between neural and non-neural elements. (141 words)

---

The emergence of cognitive science in the second half of the twentieth century offered a broad theoretical framework for understanding cognition. While its initial focus on abstract formal descriptions shifted to connectionist approaches based on neural models of cognitive architecture (Bermúdez, 2014; Haugeland, 1995; Thagard, 2013), standard cognitive science shares a fundamental “locational” commitment: whether mental processes are best seen as abstract formal processes (though exclusively realized in the brain), or as activation patterns in neural networks, they unfold in the brains of cognizers and can be described in abstraction from the body (Clark, 2008; Rowlands, 2004; Shapiro, 2012). Thus, the view is that while the organism’s body and sensorimotor systems deliver sensory input and enable behavioral output, they do not shape cognitive processing in any significant and interesting way.

In contrast, the relatively recent research program “embodied cognition” (EC) opposes the “locational” commitment, holding that at least some cognitive processes are best comprehended in terms of a dynamic interaction of bodily (non-neural) and neural processes (Foglia & Wilson, 2013). That said, EC is not a unified area of research, and the various research projects usually subsumed under the EC label lack homogeneity, established definitions (M. Wilson, 2002), and clarity about whether EC is conceived as complementing or providing an alternative to standard cognitive science.

Central claims of EC are based on findings in several disciplines, including psychology, robotics, and neuroscience (see (Barsalou, 2010). To mention a few instructive examples, researchers have demonstrated that sensorimotor variables can influence cognitive tasks (Barsalou, 2008; Hegarty, 2004; Rubin, 2006; M. Wilson & Knoblich, 2005; Zwaan, 2004), that gesturing can support the comprehension of number concepts and calculations (Andres, Olivier, & Badets, 2008; Goldin-Meadow, Nusbaum, Kelly, & Wagner, 2001; Goldin-Meadow & Wagner, 2005; Sweetser & Goldin-Meadow, 2004), and that some higher-level cognition is founded on modal systems (Martin, 2007; Pulvermuller, 2005), sometimes involving the activation of motor system simulations (Rizzolatti & Craighero, 2004).

The results of relevant research have given rise to different EC accounts that all reject the “locational” commitment, but adopt more or less radical formulations of the basic idea (see Shapiro 2012). While this is not the place to identify fine-grained differences between them, according to what we could call a *weak formulation of the EC hypothesis*, the body exerts a significant and often unexpected influence on cognitive processing, to the extent that failing to include sensorimotor aspects leads to accounts of cognition that are at best incomplete. In contrast, the *strong formulation of the EC hypothesis* states that the body acts as a (partial) realizer of cognitive processing that is distributed across neural and non-neural entities.^1^

What we propose in this paper is consistent with the weak formulation of the EC hypothesis. We draw attention to respiration and respiratory muscle activity, thus a constantly active and unique bodily and neural process that has not received attention in the EC literature. The absence of systematic interest in respiration is somewhat curious, as the characteristic rhythmic contraction and relaxation of the body that gives rise to the continual flow of air through the mouth or nose is uniquely situated at the intersection of body and mind. On the one hand, quite similar to other vital functions (e.g. digestion, endocrine and cardiovascular functions) breathing operates autonomously, adapting to environmental demands and maintaining homeostasis. On the other hand, unlike these latter functions, respiration allows conscious top-down control: not only do cognitive states modify respiratory rate and volume, but breathing can also be brought under conscious control, for example during speech production or the playing of a wind instrument.

Our paper aims to investigate respiration from an EC perspective by substantiating three hypotheses.

1. *The influence hypothesis*: besides its vital function of supplying oxygen and removing carbon dioxide, respiration exerts significant and unexpected bottom-up influence on movements, sensory perception, and cognitive processes.
2. *The mechanism hypothesis:* via respiration-driven sensory inputs, respiration influences cognition by modulating neuronal oscillations, which in turn influence the generation of action potential that realizes specific cognitive processes.
3. *The general (speculative) hypothesis:* neural synchronization through oscillations might be modulated by sensory inputs from all sensory modalities, as well as proprio- and interoceptive sensory inputs, suggesting a general mechanism through which the body influences cognitive functioning.

If (1) is true, then it is also true that a comprehensive account of cognition cannot be attained in abstraction from the respiratory activity of the body, which would lend additional support to the weak formulation of the EC. If (2) is also true, then not only is (1) further strengthened, but we have also taken initial steps towards support for the admittedly speculative (3).

To support our three theses, we draw on the emerging interdisciplinary field that investigates “neuronal oscillations” (for a review, see (Buzsaki & Draguhn, 2004), and in particular on electrophysiological studies linking oscillatory neuronal activity to the performance of sensory, motor, and cognitive tasks (Buzsaki, 2006; Buzsaki & Watson, 2012; Ward, 2003). We start with (1) offering support for the first hypothesis by providing evidence for the interaction between respiration and motor, sensory, emotional, and cognitive processes. Then, (2) we explain how neural oscillations are implicated in the synchronization of neuronal activity underlying cognitive functions and discuss evidence for a specific neuronal mechanism through which respiration influences cognitive function. Subsequently, (3) we offer support for the hypothesis that the mechanism uncovered in (2) may represent a general mechanism through which the body influences cognitive processes. Finally, in the last part of the paper (4), we work out the implications for EC, draw a parallel to the role of gesture in thinking and argue that respiration sometimes plays a double role, executing both a pragmatic and an “epistemic action” that offers a cognitively significant contribution. In such cases, the overall cognitive activity includes a loop-like interaction between neural and non-neural elements.

## 1. Support for hypothesis (1): Respiratory function beyond gas exchange

While the exchange of oxygen for carbon dioxide in the lungs is the most important physiological function of respiration, there are many known effects of breathing on motor, sensory, and cognitive functions that cannot be explained by the physiology of gas exchange. In the following we describe examples for each category. Before doing so, we find it useful to point out how the research described here differs from much earlier studies that have investigated links between cardiovascular activity with respiration and with slow oscillations of brain activity measured with EEG (Birbaumer, Elbert, Canavan, & Rockstroh, 1990). Those studies were conducted at a time where high frequency oscillations were not yet considered as a functionally relevant brain activity patterns and the changes in brain activity measured in those studies are often linked to states of sleep or arousal (Dworkin & Dworkin, 2004). There is a known influence of respiration on heart-rate and heart-rate variability (respiratory sinus arrhythmia) (Berntson, Cacioppo, & Quigley, 1993) which indirectly links breathing to blood flow changes through the brain as well as blood pressure changes which are signaled to the brain by baroreceptors and will result in changes in brain activity (Rau, Pauli, Brody, Elbert, & Birbaumer, 1993). Those forms of respiratory influence on brain activity may affect cognitive functions, but they would do so on a time scale slower than the one we are considering. Moreover, most central to our argument, those changes in brain activity are not directly linked to voluntary body movements. Our argument centers on the role of direct sensory feedback from movements of the body and we are basing it on findings linking sensory feedback from respiration to instantaneous changes in high frequency brain activity that has been widely linked to respiration.

### 1.1 The interaction between respiration and movement

One of the most basic parameters of motor control, the amount of force a muscle generates when contracting has been shown to highly correlate with the respiratory cycle. Li and colleagues measured peak handgrip force together with spontaneous and forced respiratory movements and showed that the peak force was significantly higher during forced expiration than during forced inspiration (Li & Laskin, 2006). This finding provides quantitative support for the traditional teachings in martial arts, where punches are delivered during forced expiration in order to maximize force of the punch.

Such correlation with the respiratory cycle is further supported by studies on the movements of non-respiration-related body parts, such as eye or finger movements. Eye movements have been shown to be phase-locked to respiration during sleep (Rittweger & Popel, 1998) as well as in the awake state (Rassler & Raabe, 2003). Moreover, response latency, tracking-precision and movement duration of finger movements performed to track a visual target showed significant respiratory-phase-dependent differences and that the respiratory-phase-dependence of finger movement parameters was different for finger flexion and extension movements (Rassler, 2000; Rassler, Ebert, Waurick, & Junghans, 1996). For example, Rassler and colleagues (2000) asked subjects to track a square-wave signal presented on a screen with a cursor that was moved by finger extension or flexion. Tracking by finger flexion turned out to be least precise when the movements started in the late expiration phase compared to other phases of breathing cycle. On the contrary, tracking by finger extension movements was most precise during late expiration and less precise during all other phases (Rassler, 2000).

Similar effects were shown in studies on the relation between the respiratory cycle and finger movements of pianists. Ebert and colleagues asked pianists to perform a simple finger exercise, notated in quarter notes and presented in different types of meters (e.g. 3/4 or 4/4), where the meter defines how notes are grouped (groups of 3 in the 3/4 and groups of 4 in the 4/4 meter). The exercise was performed at a comfortable tempo chosen by each pianist. Independent of the tempo, the respiratory rate assumed values that were integer ratios with the meter of the music. The authors also observed periods of constant phase relations between onsets of the meter and of the onset of inspiration. These findings demonstrate that the mental process of grouping a sequence of musical notes by various meters modulates the unconscious breathing rhythm (Ebert, Hefter, Binkofski, & Freund, 2002).

While more evidence is needed to establish this point, such correlation may perhaps occur due to the fact that the communication between motor cortical areas and motor neurons in the spinal cord is modulated with respiration. This is at least indicated in experiments using transcranial magnetic stimulation (TMS), a non-invasive method to activate cortical neurons with a rapidly changing magnetic field (Walsh & Cowey, 2000). When Li and Rymer activated the hand area of the motor cortex and measured the amplitude of resulting muscle twitches in finger muscles, they were able to show that twitches had a larger amplitude when the subjects inhaled or exhaled forcefully compared to normal, spontaneous breathing (Li & Rymer, 2011).

### 1.2. The influence of respiration on sensation

Studies on visual and auditory signal detection found that the threshold for signal detection is lower during exhalation compared to inhalation. In their experiments on visual perception, for instance, Flexman and colleagues presented visual stimuli at a low illumination adjusted to a threshold where participants detected the stimuli in only 50% of cases. When the authors analyzed whether the probability of stimulus detection varied with respiration phase, it turned out that these hard-to-perceive visual stimuli were detected significantly more often when presented during expiration compared to inspiration (Flexman, Demaree, & Simpson, 1974).

While these findings point to the correlation of sensitivity to signal detection with the respiratory cycle, there is now also evidence of a causal influence that flows from respiratory activity to signal detection. Two different studies investigated whether the detection of visual (Li, Park, & Borg, 2012) or auditory (Gallego, Perruchet, & Camus, 1991) stimuli varied with respiratory phase, by measuring reaction times in response to stimulus detection. Both studies found that during spontaneous breathing at rest, reaction times were not influenced by respiration. However, if subjects performed controlled breathing by following the rhythm of a metronome, reaction times were significantly increased during the expiration phase compared to inspiration. More recent research suggests, however, that spontaneous breathing at rest will modulate reaction times to sensory detection tasks if the stimuli and task are more complex. Zelano and colleagues (2016) very briefly (100 ms) flashed images of faces with emotional expressions of fear or surprise. Subjects had to quickly decide which expression was shown, and their reaction times were significantly longer when faces were presented during expiration compared to inspiration (Zelano et al., 2016).

Further supporting the causal influence, studies demonstrate that the perception of pain is modulated by respiration in a cycle-by-cycle manner and as a function of respiratory rate. Iwabe and colleagues (Iwabe, Ozaki, & Hashizume, 2014) have shown that pain is perceived as less severe if experienced during expiration compared to inspiration. Several studies reported that slow breathing reduces pain perception. For example, Zautra and colleagues investigated the role of respiratory rate on the perceived severity of a thermal pain stimulus and showed that focused slow breathing reduced the perceived pain (Zautra, Fasman, Davis, & Craig, 2010).

### 1.3. The influence of respiration on cognition and emotion

Conscious control of breathing or the directing of the focus of attention on the process of spontaneous breathing can be used to change emotional states and cognitive processes (Arch & Craske, 2006; R. P. Brown & Gerbarg, 2005; Paul, Elam, & Verhulst, 2007). This knowledge has been an integral part of the traditional practices of yogic breathing (R. P. Brown & Gerbarg, 2005; Jella & Shannahoff-Khalsa, 1993; Stancák & Kuna, 1994), but simple forms of controlled slow breathing, known as “combat tactical breathing” are also used by the military and special forces to reduce stress and regain focus in extremely stressful situations (Grossman & Christensen, 2011). This exercise requires counting to four during each phase of the respiratory cycle, including deliberate pauses before inhaling and exhaling.

Recent work by Zelano and colleagues suggests that the processing of fearful facial expressions and the ability to recall memorized information is modulated by respiration (Zelano et al., 2016). Interestingly, this modulation depended on subjects breathing through the nose. Photographs of faces showing expressions of either fear or surprise were shown to subjects for a very short amount of time (100 ms), and then the subjects had to quickly decide which expression the face showed. Respiration was monitored throughout the experiment. Subjects would detect fearful faces more quickly during nasal inspiration than expiration. Respiratory phase did not influence the recognition of surprised faces. When subjects were breathing through the mouth, reaction times to both types of faces increased significantly, but there was no longer a difference in detection time between inhalation and exhalation. This suggests that nasal respiration plays a special role, possibly because it involves activation of the olfactory bulb. We will discuss this in detail in the section on mechanism.

In a second experiment, Zelano and colleagues presented subjects with 180 pictures of as many different everyday objects, such as household items, animals, tools etc., and asked them to remember them. In a memory retrieval session, subjects were presented pictures from the original set plus pictures from a set of 180 objects they had not seen before. Respiration was monitored during all phases of the experiment together with the times pictures were presented. Results show that retrieval accuracy was significantly higher for images presented during the inspiration phase in the retrieval session (Zelano et al., 2016). As with the recognition of facial expressions, respiratory modulation of memory only occurred when subjects were breathing through the nose.

Taken together, the studies discussed in this section reveal how respiration correlates with motor, perceptual, emotional, and cognitive processes, which have no relevance for the primary gas-exchange function of respiration. More precisely, they provide evidence for a robust correlation between movement and the respiratory cycle, but also for *direct causal influence* on perception, emotion regulation, and cognitive processes. On such background, the next sections will discuss evidence for a neuronal mechanism, which we hypothesize underlies the respiratory influence on some cognitive processes.

## 2. Support for hypothesis (2): Respiratory modulation of neuronal activity in cognition

While an all-encompassing definition of what constitutes cognition is still missing, modern neuroscience has taken a pragmatic approach by investigating quantifiable brain functions associated with cognitive functions, including sensory perception, memory, attention, decision making, problem solving, and language processing. Studies consistently show that the occurrence of specific oscillatory patterns is highly correlated with the performance of specific cognitive tasks. Particularly, oscillations of neocortical activity in the gamma (30–100 Hz) frequency range, have been tightly linked to attention (Fries, Reynolds, Rorie, & Desimone, 2001; Gross et al., 2004; Tallon-Baudry, 2004), sensory perception (Cardin et al., 2009; Engel, Fries, & Singer, 2001; Gross, Schnitzler, Timmermann, & Ploner, 2007), decision making (Beshel, Kopell, & Kay, 2007; Siegel, Engel, & Donner, 2011), problem solving (Sheth, Sandkuhler, & Bhattacharya, 2009), memory formation (Osipova et al., 2006; Sederberg et al., 2007; van Vugt, Schulze-Bonhage, Litt, Brandt, & Kahana, 2010) and language processing (Babajani-Feremi et al., 2014; Crone et al., 2001; Towle et al., 2008).

Whereas early studies of the role of neuronal oscillations underlying cognitive functions were purely correlational, advances in experimental methods, such as new non-invasive stimulation techniques in humans and selective optogenetic manipulations of oscillatory neural networks in rodents, have provided early evidence for a causal relationship between the occurrence of cortical oscillations and sensory perception in mice (Cardin et al., 2009) and between cortical oscillations and sensory perception and working memory in humans reviewed in (Thut, Schyns, & Gross, 2011). Below we briefly discuss one example each from mouse and human studies.

Cardin and colleagues took advantage of modern optogenetic tools that allows the activation of specific types of neurons by shining light on them (Adamantidis et al., 2015). Gamma oscillations require synchronized rhythmic activation of inhibitory neurons. Using optogenetic tools to rhythmically activate inhibitory interneurons Cardin et al. were able to induce 40 Hz gamma oscillations in the whisker barrel cortex in mice, i.e. the part of neocortex that processes sensory information from the mystacial vibrissae (Brecht, 2007). By timing the mechanical stimulation of a whisker to different phases of the induced gamma oscillation, they could show that sensory input elicited the strongest response (i.e. the largest number of spikes in barrel cortex neurons) when the stimulus occurred during the negative phase of the LFP gamma oscillation (Cardin et al., 2009). Thus, sensory events coinciding with this phase of the gamma cycle have the highest probability of eliciting spikes in a large group of barrel cortex output neurons, whose highly synchronized firing would in turn maximize their ability to activate downstream structures. This type of oscillation induced neuronal synchrony or “temporal binding” has been proposed to be the neuronal mechanism behind sensory awareness and stimulus feature binding (Engel & Singer, 2001).

Non-invasive manipulation of oscillatory neuronal activity in humans can be accomplished with transcranial magnetic stimulation (TMS) techniques (Thut, Veniero, et al., 2011). Chanes and colleagues used TMS in humans to induce neuronal oscillations at high beta (30 Hz) or gamma (50 Hz) frequencies in the frontal eye field, an area involved in visual attention. The goal of the experiments was to determine a possible causal link between oscillations in these two frequency bands and visual perception (Chanes, Quentin, Tallon-Baudry, & Valero-Cabre, 2013). The results show that inducing high-beta oscillations in the frontal eye field enhanced perceptual sensitivity, allowing the subject to detect weak visual stimuli more reliably than without stimulation. The induction of gamma oscillations on the other hand did not affect perceptual sensitivity but induced a bias in individuals to give positive feedback, i.e. and increased tendency to report that a specific stimulus was perceived even if not present (Chanes et al., 2013). However, conclusions need to be qualified based on the fact that the experiments do not replicate natural neuronal oscillations. The main differences are that the experimental technique drives the networks to oscillate by external influence, following an external timing and the technique synchronizes a network defined by the reach of the magnetic field and not be the functional properties of the neurons. If one takes these caveats into account, the results are nonetheless consistent with currently held views that frequency bands of neuronal oscillations serve different functions.

Thus, in addition to the long-standing correlational evidence for a link between neuronal oscillations and cognitive function there is now also causal evidence showing that oscillations frequency-band specific effects on cognitive function. This link is essential for our hypothesis of a neuronal mechanism through which respiration exerts an influence on cognitive processes.

### 2.1. Respiration influences neuronal oscillations

Our hypothesis is in part based on the groundbreaking finding that respiration, via sensory inputs from the olfactory bulb, modulates neuronal oscillations in the delta and gamma frequency bands in the neocortex of awake mice (Ito et al., 2014). This influence of olfactory bulb activity on oscillatory neuronal activity phase-locked to respiration has long been known to occur in the brain areas involved in the processing of olfactory information (i.e. the piriform cortex) (Fontanini & Bower, 2005; Fontanini, Spano, & Bower, 2003). The olfactory bulb generates rhythmic respiration-locked neuronal activity that is caused by the rhythmic flow of air across the olfactory epithelium even in the absence of any odor (Adrian, 1951). Odors are not required for olfactory sensory neurons to fire in phase with respiration because olfactory sensory neurons have mechanosensitive properties and respond to changes in pressure caused by the rhythmic airflow through the nose (Grosmaitre, Santarelli, Tan, Luo, & Ma, 2007).

The crucial novelty in the findings by Ito and colleagues is that respiration-locked delta oscillations are present in an area of neocortex not involved in olfactory processing and that the power of gamma oscillations in that area (the whisker barrel cortex) is modulated in phase with respiration (Ito et al., 2014). The authors showed in a series of experiments that the influence of respiration on cortical oscillatory activity was almost entirely caused by respiration-locked sensory input from the olfactory bulb. Elimination of the olfactory bulb output eliminated 80-90% of the influence of respiration on cortical oscillations (Ito et al., 2014). This was the first demonstration that sensory events can modulate not only slow neuronal oscillations, but also the power of gamma band oscillations. Moreover, this can occur in a cortical area that is not involved in the processing of the (olfactory) sensory activity that caused these modulations. This is significant for our argument because the regulation of gamma oscillation power is widely implicated in cognitive functions (for recent reviews see (Bosman, Lansink, & Pennartz, 2014; Herrmann, Frund, & Lenz, 2010) and conversely, failure to regulate gamma power properly has been linked to cognitive brain disorders such as schizophrenia (e.g. (Furth, Mastwal, Wang, Buonanno, & Vullhorst, 2013) and autism (e.g. (C. Brown, Gruber, Boucher, Rippon, & Brock, 2005). Thus, the finding that respiration, via sensory inputs, can modulate gamma oscillations in areas of necortex not related to olfaction suggests that respiration can modulate neuronal processes underlying cognitive function.

The group went on to demonstrate that respiration-locked oscillations occurred in many other non-olfactory areas of the mouse neocortex (Figure 1) and, most relevant to our point here, that respiration-locked oscillations and respiration-locked modulations of gamma power also occur in the human neocortex (Heck et al., 2016; Liu et al., 2015). Human cortical recordings were obtained from pharmaco-resistant epilepsy patients who received implants of subdural grid electrodes (electro-corticogram) and showed respiration-locked modulation of gamma-power in the frontal, parietal, and temporal neocortex (Heck et al., 2016; Liu et al., 2015). Those areas, particularly the frontal cortex, are association areas essential for cognitive function. These findings received support by a later study, also conducted in epilepsy patients, showing respiration-locked modulations of the power of delta, theta, and beta oscillations in the piriform (olfactory) cortex, the amygdala, and the hippocampus, when patients were breathing through the nose (Zelano et al., 2016). The findings support the general view that olfactory sensory input can cause a modulation of higher frequency oscillations.

**Figure 1:**
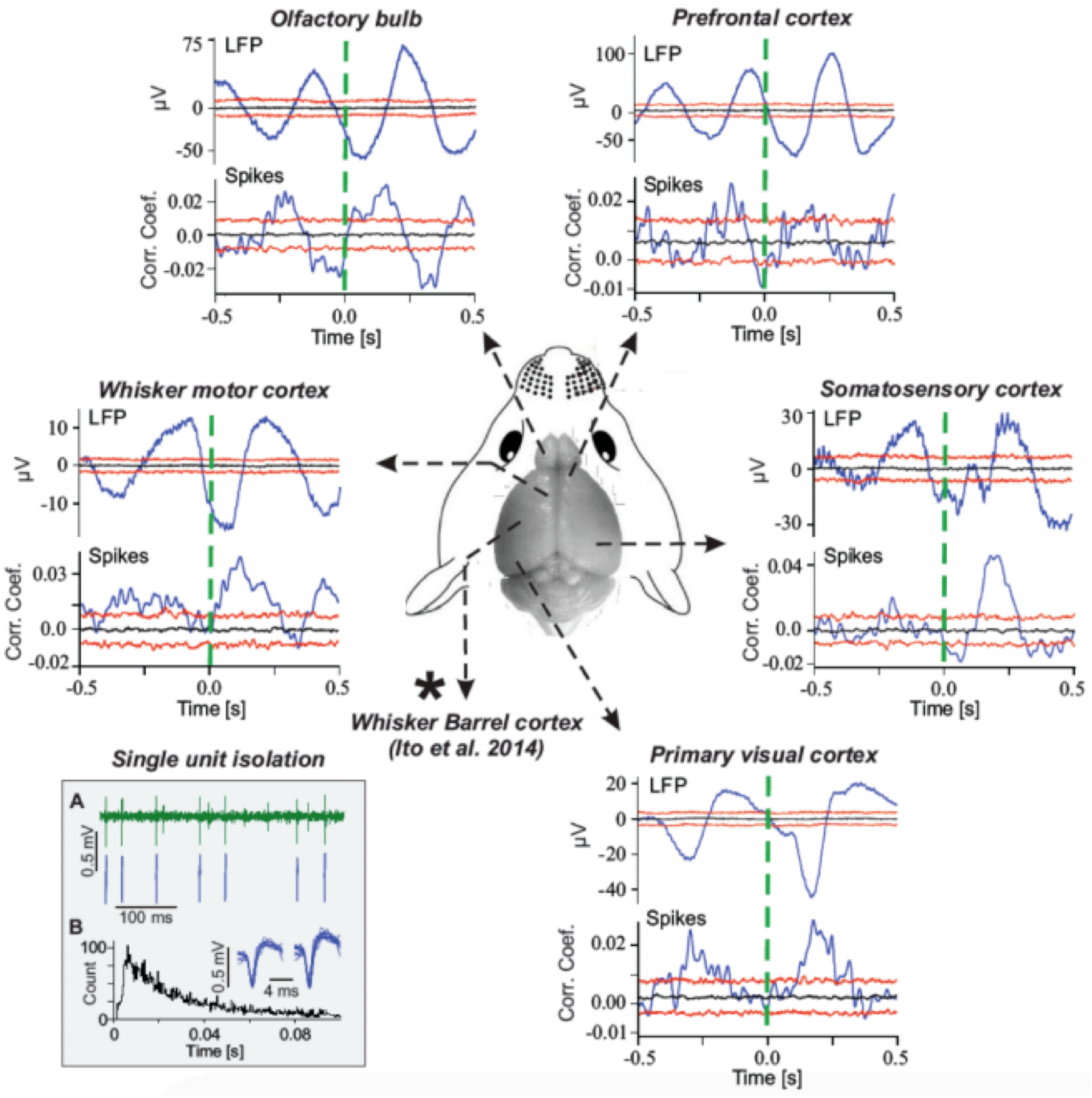
Spike and LFP activity in different areas of the neocortex of the awake mouse are rhythmically modulated in phase with respiration. Blue lines in upper histograms are averaged LFP signals, aligned on the time of end of expiration (dashed vertical green lines). Blue lines in lower histograms are cross-correlograms of spike activity with the end-of-expiration times. Horizontal black and red lines in each histogram represent the predicted median (black) and the 5 and 95 percentile boundaries (red) of the surrogate data obtained by random shuffling (bootstrap statistics). LFP averages and spike-respiration cross-correlation functions exceeding the red boundaries are considered significant (p > 0.05). Square insert at the bottom left shows an example of a single unit recording in primary motor cortex. (A) In insert: raw spike recording with extracted spike waves of sorted single unit spikes shown below. (B) In insert: inter-spike interval (ISI) histogram showing the typical shape of a single unit with effects of refractory period. Traces above ISI show overlays of the first and last 10 spikes recorded over 300 seconds (Figure from Heck et al.). 2016).

It is worthwhile at this point to clarify that the observed modulation of gamma activity in form of a sudden increase in gamma amplitude has two possible causes: one is an increase in power of an ongoing low-amplitude oscillation or the onset of a gamma burst in a network that, in the time immediately preceding the observed gamma oscillation, did not produce any oscillation in that frequency band (Buzsaki & Wang, 2012). The validity of our argument does not depend on one or the other mechanism to be involved. Simulation experiments with graph theory models have replicated respiration-locked modulation of gamma power (Heck et al., 2017; Heck et al., 2016). Additional studies are needed to determine whether respiration-locked activity can influence the initiation of gamma bursts. Support for such a mechanism comes from a recent study in mice, showing that the initiation of hippocampal sharp wave ripples (a high-frequency network event linked to memory function) is phase locked to respiration (Liu, McAfee, & Heck, 2017).

The experiments in mice (Ito et al., 2014) showed a residual influence of respiration on cortical activity after the olfactory bulb was removed. This indicates that other respiration-locked inputs also contribute to respiratory modulation of brain activity. There are two main sources to consider: other sensory inputs—proprioceptive and interoceptive—, and respiration locked activity reaching higher brain areas via ascending projections from the spinal cord. Ascending respiration-locked activity originating from the brain stem respiratory circuits (Feldman & Del Negro, 2006) would provide respiration-locked activity independent from sensory consequences of respiration. While such projections would likely contribute to respiration-locked oscillations in higher brain areas, their relative importance compared to the numerous sensory inputs remains to be determined. Existing data in mice and humans currently suggest that the bulk of the effect depends on the olfactory bulb and not on brain stem signal.

Respiration causes both conscious and unconscious sensory inputs. Consciously accessible sensations of normal, unobstructed breathing include odor perception, the mechanical and thermal sensation of air flowing through nose, mouth, and upper airways, the auditory component of heavy breathing after a workout, and the proprioception of movements of the chest and abdomen. Unconscious respiration-locked sensory signals include, of course, signals from chemosensors in the vascular system, which signal breath-by-breath fluctuations of CO_2_ and O_2_ levels in the blood but also interoceptive signals from receptors in the lungs and diaphragm and also internal organs, which are moved with every breath, particularly during abdominal breathing. Provided that the olfactory sense is much more prominent in mice than in humans, it is likely that the influence of proprio- and interoceptive respiration-locked sensory inputs on brain activity are relatively larger in humans than in mice. The findings by Ito et al. that removal of the olfactory bulb reduces respiration-locked cortical activity by 80-90% supports the notion that respiration-locked brain activity in mice is dominated by inputs from the olfactory bulb (Ito et al., 2014).

However, the clearly important role of olfactory bulb in mice as well as in humans should lead us to think that individuals with non-congenital anosmia display cognitive deficits. ^2^ In mice and rats, removal of the olfactory bulb does in fact have dramatic effects on behavior and is the established experimental model for inducing depression in rodents (Song & Leonard, 2005). Since the olfactory bulb in humans is smaller relative to the volume of the rest of the brain, the cognitive effects of anosmia are expected to be correspondingly smaller in humans than in mice. This is indeed the case. In fact, patient studies show that the degree of depression scales with the severity of anosmia (Kohli, Soler, Nguyen, Muus, & Schlosser, 2016). Moreover, odor identification has a strong relationship with memory performance in healthy older adults and is a significant predictor of future cognitive decline (R. S. Wilson et al., 2007).

Overall, experiments in mice and humans show that respiration modulates neuronal activity at the frequency of respiration and that it modulates the power of higher frequency oscillations, including gamma oscillations. Furthermore, the data thus far suggest that the strongest driving force behind this effect is respiration-locked sensory input to the cortex from the olfactory system. While it is relatively straightforward to understand how neuronal activity in the neocortex can become synchronized to a rhythmic sensory input, it is less clear how an external input can modulate the power of intrinsic, higher frequency cortical oscillations. A recent modeling study replicated the power modulating effect of respiratory sensory inputs in a graph theoretical model that mimicked basic features of neocortical excitation/inhibition balance and neuronal connectivity, thus suggesting that the phenomenon requires no assumptions beyond the intrinsic feature of cortex-like network architectures (Heck et al., 2017).

In support, we may note that the respiratory influence on brain activity is largely consistent with mammalian brain evolution. X-ray studies of fossil skulls of early Jurassic mammaliaforms suggest that the increase in olfactory bulb size was a crucial event in the evolution of the mammalian brain (Rowe, Macrini, & Luo, 2011). Consequently, early mammals likely had an excellent sense of smell, which greatly improved their ability to live their predominantly nocturnal lifestyles and allowed them to evade predation by diurnal carnivores. Provided what we now know about the influence of respiration-locked olfactory bulb activity on neuronal oscillations in rodent brains, it is likely that respiration created an ever-present fundamental neuronal rhythm that may have shaped the temporal organization of neuronal activity during mammalian brain evolution. Some highly preserved spatiotemporal patterns of neuronal activity such as default mode networks (Greicius, Supekar, Menon, & Dougherty, 2009) might thus—in ways we do not yet fully understand—be the result of the incessant interaction between intrinsic network activity and respiration-locked neuronal rhythms.

### 2.2. Neural oscillations influence action potential generation

Having established that respiration modulates neuronal oscillations in the neocortex, it is necessary to show how these oscillations, which are typically measured as local field potentials (LFPs) or electroencephalograms (EEGs), do influence neuronal output in the form of action potentials (also “called spikes”). Action potentials or spikes are transient (1 ms) all-or-nothing electrical signals that are the long-distance carriers of precisely timed information between connected neurons (Schuetze, 1983). Neuronal communication without action potentials also exists but does not provide long distance communication nor temporal precision: Neighboring neurons can exchange graded voltage information through gap-junctions formed where their membranes touch (Dermietzel & Spray, 1993), and there are also specialized cells which show graded transmitter release (Heidelberger, 2007). However, the most common form of neuronal communication is via action potentials propagated along axons.

Oscillations of EEG or LFP activity reflect mostly synaptic activity and thus the rhythmic modulation of neuronal *excitability*, which is a measure of the likelihood that a given neuron will generate an action potential in response to synaptic inputs. Periods of increased excitability represent a “window of opportunity,” during which a neuron is more likely to respond with spiking activity to synaptic inputs.

That neuronal action potential firing in the cortex is synchronized to oscillations in the LFP signal has been demonstrated in several experimental settings and cortical areas where LFPs and spikes were measured simultaneously (Eckhorn & Obermueller, 1993; Gray, Kînig, Engel, & Singer, 1989; Jacobs, Kahana, Ekstrom, & Fried, 2007; Murthy & Fetz, 1996; Nase, Singer, Monyer, & Engel, 2003). Spikes typically coincide with the negative wave of LFP oscillations, which correspond to the synaptic influx of positively charged ions into the dendrite of a cell. The exact shape of LFPs depends on the position of the recording electrode tip relative to the cell body (Buzsaki, Anastassiou, & Koch, 2012) but the negative phase recorded at the level of the dendrite generally reflects increased excitatory synaptic input and thus increased probability of spike activity in the postsynaptic cells (Buzsaki et al., 2012). Oscillations of LFPs thus organize spike activity (or more precisely, spike probability) in time (Figure 2). One theory is that neuronal oscillations serve the purpose to temporarily synchronize spike activity in different parts of the brain, as a means to enhance neuronal communication between task-related parts of the brain for the duration of the task performance (Fries, 2005, 2015; Womelsdorf et al., 2007).

**Figure 2:**
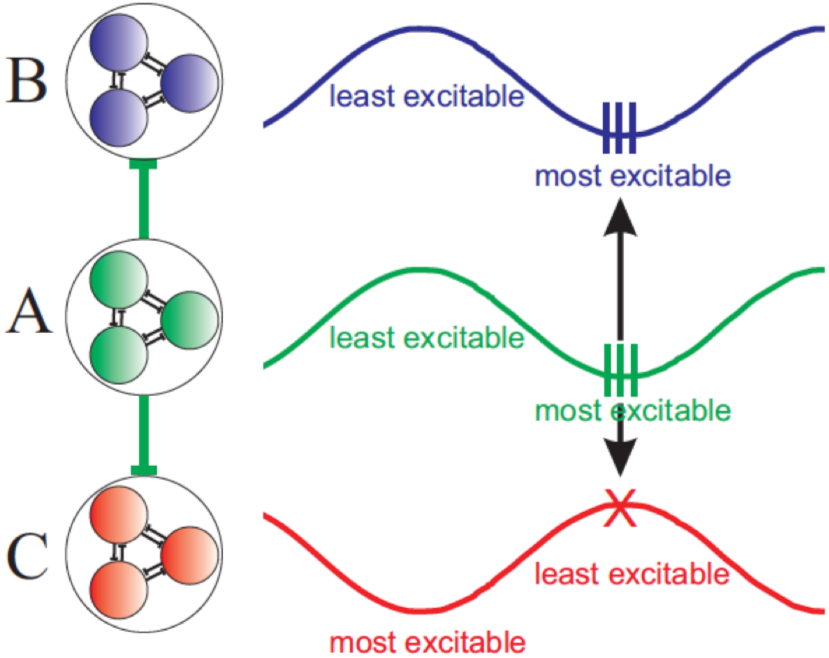
Graphical illustration of how the phase relationships between LFP oscillations in different brain areas (A, B, C) can control neuronal communication and spike synchronization. Neurons in area A send excitatory projections to areas B and C. The phases of LFP oscillations in A and B are ideally aligned in such a way that neurons in both areas simultaneously reach a state of high excitability. Consequently, there is a high probability that neurons in area B will generate spikes in response to incoming spikes fired by neurons in area A. By contrast, LFP oscillations between A and C are in anti-phase, so that A’s action potentials reach C at the time when C is least excitable, minimizing the probability of signal propagation. The above example shows the two extremes of LFP phase alignments. Any phase in between the two alignments will lead to intermediate spike propagation probabilities.

### 2.3. Action potential realizes cognitive processes

All aspects of mammalian brain function, from motor control to complex cognitive processes, require that neurons generate or “fire” action potentials in response to synaptic inputs. This can be shown experimentally by selectively preventing neurons from generating spikes while leaving all other aspects of neuronal function intact. Tetrodotoxin (TTX) is a neurotoxin found in puffer fish that prevents neurons from generating action potentials by selectively blocking the function of voltage dependent sodium channels, which are essential for action potential generation and propagation (Narahashi, 2008). Experimental proof for the fact that spike generation is essential for brain function including cognitive function comes from countless experiments using small amounts of locally applied TTX to block spike firing in specific brains and then demonstrating that the function of that brain structure is eliminated (e.g. (McLaughlin & See, 2003; Wesierska, Dockery, & Fenton, 2005).

How are cognitive processes realized as spike activity? There is currently no widely accepted answer to this question. However, electrophysiological and theoretical studies over the past decades have provided substantial evidence that information processing in the brain is based on the synchronous firing of action potentials in large groups of neurons (Diesmann, Gewaltig, & Aertsen, 1999; Uhlhaas et al., 2009). In the following, we discuss why synchronous spike firing is the most reliable way for the representation and transmission of information in neural networks and present two prominent theories of cognitive function based on synchronized population activity.

The most basic argument for the importance of synchronized spiking for cognitive function is that neurons generate action potentials most reliably and with the highest temporal precision in response to synchronized synaptic inputs (Abeles, 1991; Koenig, Engel, & Singer, 1996; Rodriguez, Aertsen, & Heck, 2007).

There is experimental evidence for a link between cognitive brain functions and synchronized spike activity. For example, prefrontal cortical neurons in primates synchronize their spike firing during the performance of a task requiring a context-dependent decision to either respond to a signal with a movement (“go” condition) or to ignore the signal (“no-go” condition) (Vaadia et al., 1995)..

In a more recent study, Svoboda and colleagues (Komiyama et al., 2010) describe the learning-related formation of synchronously active groups of neurons (“cell assemblies” in Donald Hebb’s terminology, (Hebb, 1949) in the tongue motor cortex of mice, who learned to lick a water reward in response to the presence of a specific odor and to not lick in response to a different odor. These authors suggested that the formation of cell assemblies is a signature of learning-related cortical network plasticity (Komiyama et al., 2010).

### 2.4 Respiratory modulation of brain oscillations does not depend on blood-oxygen fluctuations

It is important to emphasize that the influence of respiration on brain oscillatory activity does not depend on rhythmic fluctuations of blood oxygen levels. The most comprehensive evidence for the role of sensory inputs in driving respiration-locked oscillations and modulations of power in higher frequencies comes from the work of Ito et al. (Ito et al., 2014) in mice. These authors performed 3 control experiments which all supported the notion that respiration locked sensory input from the olfactory bulb was the main driving force behind respiratory modulation of brain oscillations. First, removing the olfactory bulb eliminated the influence of respiration on oscillations in the somatosensory cortex. Second, in a tracheotomized mouse under anesthesia natural breathing draws air into the lungs through the open trachea, bypassing the olfactory sense. Under these conditions respiration did not modulate cortical neuronal oscillations. However, when air was rhythmically pumped through the nose at a rate and volume resembling airflow during natural breathing, cortical oscillations were modulated in phase with the artificial airflow, not with the rhythm of the oxygen-providing natural breathing through the open trachea. Finally, the authors applied electrical stimuli to the olfactory bulb and could show that cortical neuronal activity could be influenced by electrical stimulation of the bulb.

In humans, Zelano and colleagues found that switching from nose-to mouth breathing reduced the effect of respiration on cortical oscillations and the effect of respiratory phase on emotional responses and memory performance (Zelano et al., 2016). Again, if oxygen fluctuations were the primary driving force it would not matter how air gets into the lungs. The fact that nasal breathing had a stronger influence on cortical oscillations in humans supports our hypothesis that the effect is due to respiration locked modulation of sensory inputs. We cannot entirely exclude an effect of blood oxygen level fluctuations on cortical rhythms but it seems to be small compared to the effects related to sensory inputs.

To sum up, in order to substantiate our claim of a neuronal mechanism through which the influence of respiratory activity influences cognitive processes, we started out by looking closer at the link between neuronal oscillations and cognitive functions. First, it was shown that respiration modulates both the neuronal activity at the frequency of respiration and the power of higher frequency oscillations. In a second step, we demonstrated that oscillations influence and synchronize neuronal output in the form of action potentials. In a third step, we presented two prominent theories of cognitive function based on synchronized population activity showing that cognitive processes are realized through action potentials. To be clear, there is no claim that oscillations are identical with, co-extensive with, or somehow amount to mental processes. We argued that respiration – via sensory inputs – influences neuronal oscillations, which in turn influences action potential generation that realizes cognitive processes. We think that the abundance of reports of correlations between gamma oscillations and cognitive performance in the entire range of brain areas thus far investigated supports the causal claim in animal models and humans (Cardin et al., 2009; Thut, Schyns, et al., 2011).

## 3. Support for hypothesis (3): sensory inputs to the neocortex modulate cognitive processes by modulating neuronal oscillations

We have thus far focused on the special case of respiration as a modulator of cognitive function and pointed to olfactory sensory input as the underlying main cause of respiratory modulation of neuronal oscillatory activity. However, it may be reasonable to assume that the spike activity representing olfactory sensory input are no different from spike trains representing sensory inputs from other sources, such as other primary senses as well as proprio- and interoceptive sensations. Since evidence for the modulation of cortical oscillations by sensory inputs has only been introduced very recently (Ito et al., 2014), there is not much experimental evidence from other modalities. However, we know of at least one example from a study of a study of saccadic eye movements in a non-human primate (Ito, Maldonado, & Grun, 2013). Ito and colleagues identified saccade-related changes in the power and phase relationships of LFP oscillations in the primary visual cortex of freely viewing capuchin monkeys. These changes occurred in a broad range of frequencies, with the phase of delta-theta oscillations found to be entrained to the rhythm of repetitive saccades and an increase in power of alpha-beta and low-gamma band oscillations time-locked to the onset of saccades. Thus, the change in visual sensory input associated with a saccadic eye movement causes - amongst other effects - also an increase in the power of neuronal oscillations, including gamma band oscillations.

On such basis, our admittedly speculative hypothesis is that all sensory inputs to the neocortex exert differing degrees of control over cortical oscillatory activity and that this mechanism represents a possible neuronal basis of embodied cognition. While the examples we have provided thus far were limited to sensory inputs from either non-moving body parts (nasal air flow) or from eye movements, which are not typically at the center of considerations of embodied cognitive processes, we would like to once again discuss gesture, partly because of its particular place in the EC literature. We have earlier suggested that respiration operates somewhat analogously to gesture in terms of its cognitive function. We now propose that respiration functions analogously in terms of the underlying neural mechanism.

A study of the effects of observed gestures on brain activity by Biau and colleagues measured oscillatory brain activity in subjects watching a speaker who used hand gestures for emphasis and while watching the speaker giving the same speech but without any gestures. The authors showed that the use of hand gestures by the speaker caused an increase in phase-locking or synchronization of theta oscillations over the left fronto-temporal scalp region (Biau, Torralba, Fuentemilla, de Diego Balaguer, & Soto-Faraco, 2015). While this shows that observed gestures do indeed modulate neuronal oscillations and synchronization related to cognitive processes, this result does not allow conclusions about the effect of the same gestures on the speaker’s brain, which is the subject of our argument. After all, the viewer perceives the gesture visually while the speaker would receive proprioceptive sensory inputs. An experiment designed to directly test our hypothesis would have to measure oscillatory neuronal activity of the speaker’s brain while the speech is given with and without gestures. To the best of our knowledge, such a study has not yet been conducted.

There are, however, electrophysiological studies of the motor system that can provide valuable clues as to the potential effect of a speaker’s gestures on oscillatory brain activity. Numerous studies have investigated of oscillatory neuronal activity in the sensorimotor cortical areas of primates and humans during limb-movement with the goal of understanding the neuronal mechanisms of motor control. After oscillations were discovered as an omnipresent part of cortical neuronal activity, their role in motor control became a focus. Magnetoencephalographic (MEG) recordings of cortical activity in humans performing simple, single-joint limb movements show that these movements induced increases in neuronal gamma oscillations in sensorimotor areas of neocortex (Cheyne, Bells, Ferrari, Gaetz, & Bostan, 2008). One would, of course, like to know how the performance of more natural multi-joint movements, or ideally gestures performed to emphasize speech, modulates brain activity, but MEG measurements require the head to remain perfectly still, and EEG is extremely sensitive to electrical artifacts caused by muscle activation when limbs are moved. The closest we can currently get to finding out how more natural movements modulate oscillatory brain activity is by using micro-electrodes to record neuronal activity directly inside the brain tissue in behaving primates. Recordings of spike and oscillatory LFP activity have been performed in the primary motor cortex of monkeys while they performed voluntary hand movements. The results showed that at the moment of movement-onset oscillatory and spike activity significantly changed in complex ways (Donoghue, Sanes, Hatsopoulos, & Ga†l, 1998).

While this study did not reach a conclusion as to the functional significance of these changes, other findings suggest that oscillations in the motor system improve the transmission of spike signals from the cortex to the spinal cord by synchronizing activity in cortical projection- and spinal motor neurons, a phenomenon termed “corticomuscular coherence” (Salenius & Hari, 2003). Studies in humans using MEG have shown that the synchrony of oscillations in the sensorimotor cortices is highly modulated during a motor task requiring the subject to control position of a handle with precision finger grip (Kilner et al., 2003). Oscillations were highly synchronized during position-holding and desynchronized when the task required the subject to move the handle to a new position. When the new position was reached, synchronization resumed (Kilner et al., 2003). This latter study shows that cortical oscillations in humans are modulated by movements of extremities. This and other similar studies focused on motor control or observation of movements and focused on the neuronal activity in sensorimotor cortical areas. Thus, we currently don’t know if movements or observations of movements result in changes of neuronal oscillations in cortical areas involved in cognitive tasks. That this is at least in principle possible, however, has been demonstrated by language processing studies showing that action-words, i.e., words evoking images of body movements, result in the simultaneous activation of association and motor cortical areas (Pulvermuller & Fadiga, 2010).

Additional indications of the importance of proprio- and interoceptive sensory inputs for the spatial and temporal structure of cortical oscillatory activity come from an investigation of the effects of spinal injury on brain activity. Aguilar measured neuronal activity in the forepaw and hindpaw somatosensory cortex of anesthetized rats before and after complete thoracic transection of the spinal cord (Aguilar et al., 2010). The resulting loss of proprio- and interoceptive sensation below the level of the transection resulted in immediate changes in large-scale cortical activity patterns, which included a significant reduction of spontaneous activity and the onset of a slow wave oscillation that was synchronized across large areas of cortex (Aguilar et al., 2010). Thus, the presence of proprio-interoceptive sensory inputs clearly shape cortical neuronal activity. The link between the sensory inputs caused by, e.g., hand and arm gestures for cortical activity is currently unkown, but can principally be addressed experimentally.

## 4. The dual function of respiration

Our investigation of respiration demonstrates that the organism’s body and sensorimotor systems not only deliver sensory input and enable behavioral output, but can also shape cognitive processing in a significant and interesting way. In other words, our findings supports the weak formulation of the EC hypothesis, showing that the body may exert a significant influence on cognitive processing, to the extent that failing to include these sensorimotor aspects would lead to incomplete accounts of cognition. Moreover, in addition to contributing to the field by adding respiration to the growing palette of processes that can be fruitfully studied from an EC perspective, we identified a specific mechanism through which cognitive processes are amenable to modulation by rhythmic respiratory activity. We did not merely focus on the manner in which the brain implements cognitive functions, but also sought to identify neural mechanisms that help explain dynamic interactions between the brain and the non-neural body. However, in part because breathing and respiratory sensory and motor activity has not received sustained attention in the in the EC literature, it is worth adding further reflections on its place in the landscape.

To achieve this, it is helpful to start by drawing a parallel to gesture, which is often used in the EC literature as a model of how the body shapes cognition (Clark, 2008; Gallagher, 2005). Much of this literature is based on the observation that gesture is more than an aid for the outward communication of previously formed thoughts: not only do we gesture in the absence of others, but congenitally blind individuals also gesture when expressing themselves verbally (Iverson & Goldin-Meadow, 1998, 2001). Indicating that gesture functions as part of cognition, instructing participants not to gesture has significant detrimental effects on cognitive performance (Goldin-Meadow et al., 2001; Sweetser & Goldin-Meadow, 2004). The alternative explanation that complying with the instruction to refrain from gesturing is adding to the load was ruled out, indicating that gesturing is more than a motor act that articulates a cognitive process in the brain. Instead, its role could be to reduce the cognitive load, which would establish gesture as a genuine part of a process that couples neural and bodily resources in a loop-like manner (see also (McNeill, 2005). Put in different terms, gesture performs two separate tasks. Using a distinction coined by David Kirsh and Paul Maglio (Kirsh & Maglio, 1994), gesture is both a “pragmatic action” (executed to performed a physical task), and an “epistemic action” (executed to reduce the cognitive load when performing a task).

In light of the example of gesture, we want to suggest that respiration operates somewhat analogously. We have in the last section speculated that respiration and gesture may function analogously in terms the underlying neuronal mechanism, i.e., sensory modulation of neuronal oscillations. Here, we want to suggest that they might function analogously, fulfilling a double role as pragmatic and epistemic action. On the one hand, respiration is clearly a “pragmatic action”: it is executed to directly accomplish the physical goals of lung ventilation and gas exchange. On the other hand, however, it is at least sometimes also an “epistemic action”: its execution provides the organism additional proprioceptive information that produces a transitory, but cognitively significant, contribution.

Evidence for the hypothesis of respiration as occasionally an epistemic action, comes from studies on changes in respiratory patterns during cognitive tasks. It is well known that healthy breathing exhibits balanced variability consisting of mainly correlated variability and some random variability. Such variability allows the respiratory network to adapt to changes in environmental demands (Wuyts, Vlemincx, Bogaerts, Van Diest, & Van den Bergh, 2011). Vlemincx and colleagues (Vlemincx, Taelman, De Peuter, Van Diest, & Van den Bergh, 2011) investigated random and correlated respiratory variability and sigh frequency during demanding mental arithmetic tasks and non-stressful attention tasks. The execution of the tasks was associated with different patterns in respiratory variability. Respiratory measures demonstrated that the respiratory patterns were attuned to cognitive tasks: stressful tasks with high mental load induce random variability, while a task involving non-stressful attention reduce total variability. Moreover, sighing resets the balance when respiration lacks variability or becomes overly irregular.

A recent fMRI study investigated the effects of respiratory fluctuations on memory, providing further evidence for the thesis that particular respiration patterns are linked to cognitive events (Huijbers et al., 2014). Using phase-locking analysis methods regularly applied in electrophysiological studies, this study focused on the relationship between respiratory fluctuations and stimulus presentation and found that the respiratory cycle phase-locks to the stimulus presentations (word stimuli), yielding a respiratory response that differs significantly from baseline. Moreover, the study showed that words that were later remembered elicited a higher-amplitude respiratory response than words that would not be remembered. This suggests a link between the amplitude of respiratory movements and successful cognitive performance, demonstrating that respiration correlates both with immediate and later memory performance. Analysis of the fMRI signals obtained during the task showed that signals related to the encoding of the word memories were strongly modulated by respiration in the posterior midline region of the brain, a region considered part of the “default mode network” (Greicius et al., 2009) involved in the performance of a variety of cognitive tasks. The authors take their findings to “indicate a region-specific interaction between respiration and the fMRI signal associated with successful task performance” (Huijbers et al., 2014).

Overall, while respiration patterns do not passively attune to different types of cognitive tasks, but figure as active elements in an overall cognitive economy, respiratory activity can be seen as part of the causal loop that underlies cognitive functioning: it is affected by cognitive states and tasks, while at the same time affects the underlying mechanisms of the relevant cognitive operations (Heck et al., 2017; Heck et al., 2016; Huijbers et al., 2014; Ito et al., 2014; Zelano et al., 2016) (see also (Ramirez, 2014). If it is true that a specific type of interaction between respiration and neural activity linked to cognition predicts successful task performance, then we may conclude that—akin to gesture—respiration sometimes plays a double role. It executes *both a pragmatic and an epistemic action* in which its adaptive performance provides the organism additional information, generating a cognitively noteworthy contribution. Moreover, much like gesture, it provides an example for how neural and bodily elements interact to create an integrated but distributed cognitive circuitry that sometimes functions in synchrony to ease the cognitive load. In such cases, the overall cognitive activity is embodied in the sense that it is best understood not as decomposing into internal neural activity, but as including a loop-like interaction between neural and non-neural elements via sensory signals that modulate cognition-relevant brain activity phase-locked to the body movements that cause the sensory signal.

### Concluding remarks

EC approaches oppose the “locational” commitment of standard cognitive science, holding that at least some cognitive processes are best comprehended in terms of a dynamic interaction of bodily (non-neural) and neural processes. The findings presented in this paper offer support for the weak EC hypothesis, according to which the body, via sensory signals, exerts a significant and unexpected influence on cognitive processing. Importantly, the nature of the influence is such that a comprehensive description of cognitive processing requires taking into account the complex interaction between bodily and neural processes.

Substantiating our first hypothesis, we demonstrated that unrelated to its gas-exchange function, respiration-related sensorimotor activity exerts a systematic bottom-up influence on motor, perceptual, emotional, and cognitive processes. In a second hypothesis, we advanced a neuronal mechanism through which the influence of respiratory activity on cognitive processes occurs. We focused on the link between neuronal oscillations and cognitive functions, and demonstrated that respiration influences neuronal oscillations, which in turn influences action potential generation that realizes cognitive processes. Thus, in the first two parts of the paper, we both added respiration to the growing palette of processes that can be fruitfully studied from an EC perspective and identified a specific mechanism through which cognitive processes are amenable to modulation by respiratory activity.

Based on the evidence for this mechanism, we presented a third, more speculative hypothesis, according to which all sensory input to the neocortex exert differing degrees of influence over cortical oscillatory activity, representing the neuronal basis of embodied cognition. While support for this hypothesis is relatively scarce, we found indications that cortical oscillations in humans and non-human primates are modulated by movements of extremities, which would support the thesis that gesture may function analogously to respiration in terms the underlying neuronal mechanism. In the last part of the paper, we situated our findings within the EC literature. We argued that respiration operates somewhat analogously to gesture in terms of its cognitive function, providing an example how neural and bodily elements interact to create a distributed cognitive circuitry that eases the cognitive load by a loop-like interaction between neural and non-neural elements. Respiration provides an example of how cognitive problem-solving is sometimes assisted by a sort of sensory-motor driven regulation that may be described as an epistemic action.

Perhaps by highlighting the parallel to gesture, some may argue that our account not only shows that respiration causally impacts cognition and helps lightens the load, but also supports the strong formulation of the EC hypothesis, according to which respiration itself forms a proper part of cognitive processes. However, at least in light of the current evidence, our account supports the weak formulation of EC, and demonstrates that a full understanding of cognitive processes must include the interactions between task-related processes distributed across brain and body and their coupling via sensory activity.

## Footnotes

1 More radical approaches maintain that in some cases, not only non-neural, but also non-bodily realizers can be seen as constitutive for cognitive processing (Clark, 2008; Clark & Chalmers, 1998).

2 We thank an anonymous referee for pointing out this issue to us.

